# A Genomic Language Model for Chimera Artifact Detection in Nanopore Direct RNA Sequencing

**DOI:** 10.1101/2024.10.23.619929

**Authors:** Yangyang Li, Ting-You Wang, Qingxiang Guo, Yanan Ren, Xiaotong Lu, Qi Cao, Rendong Yang

## Abstract

Chimera artifacts in nanopore direct RNA sequencing (dRNA-seq) can significantly distort transcriptome analyses, yet their detection and removal remain challenging due to limitations in existing basecalling models. We present DeepChopper, a genomic language model that precisely identifies and removes adapter sequences from base-called dRNA-seq long reads at single-base resolution, operating independently of raw signal or alignment information to effectively eliminate chimeric read artifacts. By removing these artifacts, DeepChopper substantially improves the accuracy of critical downstream analyses, such as transcript annotation and gene fusion detection, thereby enhancing the reliability and utility of nanopore dRNA-seq for transcriptomics research.

## Main

Long-read RNA sequencing technologies are revolutionizing transcriptomic research by providing unparalleled resolution for detecting complex splicing and gene fusion events often missed by conventional short-read RNA-seq methods. Among these technologies, Oxford Nanopore Technologies (ONT) dRNA-seq stands out by sequencing full-length RNA molecules directly, preserving native RNA modifications and allowing a more accurate and comprehensive analysis of RNA biology. This approach bypasses the inherent limitations of cDNA-based sequencing methods, such as artifacts arising from reverse transcription, template switching, and Polymerase Chain Reaction (PCR) amplification [1, 2].

Despite these advantages, a critical question remains: Does ONT dRNA-seq introduce technical artifacts? A previous study has suggested that dRNA-seq might generate chimera artifacts, leading to multi-mapped reads [3]. These artifacts may result from ligation during library preparation or chimeric reads produced by software missing the open pore signal, potentially confounding downstream analyses such as transcriptome assembly, quantification, and detection of alternative splicing and gene fusion events. Detecting these chimera artifacts is challenging because long-read aligners often produce chimeric alignments from such artifacts that are indistinguishable from those derived from true gene fusion events. Importantly, chimeric read artifacts frequently contain internal adapter sequences [3], which could theoretically serve as a distinguishing feature to differentiate them from true gene fusion-derived reads. However, ONT dRNA-seq basecallers, trained in RNA, struggle to properly call these DNA-based adapter sequences under an RNA model [4]. As a result, current adapter detection tools cannot exploit this feature to eliminate chimeric read artifacts, leaving the issue unresolved.

To address these challenges, we developed DeepChopper, a Genomic Language Model (GLM) for long-read sequence analysis. Leveraging recent advances in Large Language Model (LLM) that can interpret complex genetic patterns [5], DeepChopper processes long genomic contexts with single-nucleotide resolution. This capability enables precise identification of ONT adapter sequences within base-called long reads, facilitating the detection and removal of chimeric read artifacts in dRNA-seq data. DeepChopper analyzes individual FASTQ records to locate and process adapter sequences: it splits reads containing internal adapters into multiple records (Fig. 1a) and trims adapters at the 3′ ends, thereby preserving genuine biological sequences while eliminating chimera artifacts.

**Fig. 1.**
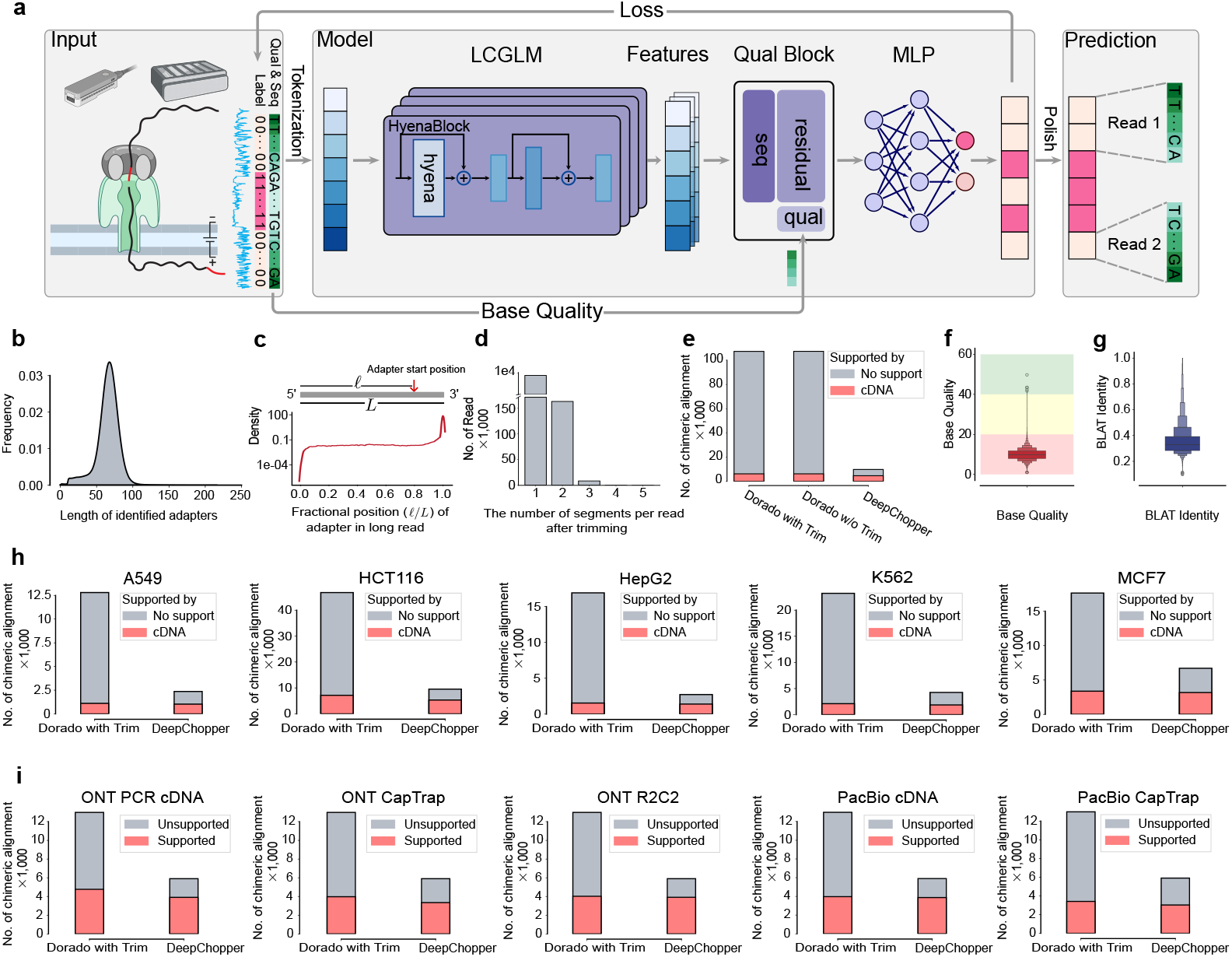
Detection of chimeric read artifacts in dRNA-seq data using DeepChopper and validation with orthogonal sequencing platforms. (a) Overview of the architecture of the DeepChopper model. Created with BioRender.com (b) Length distribution of predicted adapters by DeepChopper in VCaP dRNA-seq data. (c) Distribution of relative adapter position along read length in VCaP dRNA-seq data. Grey rectangle represents a long read from 5′ to 3′. Relative position is calculated as the ratio of the length before DeepChopper predicted adapter start position to the total read length. (d) Distribution of segments per read after trimming: 1 segment indicates 3′ end adapter trimmed, while 2 or more indicate internal adapters trimmed. (e) Chimeric alignments (in thousands) for VCaP dRNA-seq reads processed by Dorado with and without adapter trimming, and DeepChopper. DeepChopper greatly reduces chimeric alignments not supported by direct cDNA sequencing. (f) Distribution of base qualities from identified internal adapters by DeepChopper. Background colors indicate quality levels: green (high), yellow (medium), and red (low). (g) BLAST-like alignment tool (BLAT) identity distribution of the internal adapter sequences mapping against human reference genome. (h) The number of chimeric alignments (in thousands) for A549, HCT116, HepG2, K562, and MCF7 cell lines processed by Dorado with adapter trimming and DeepChopper. DeepChopper consistently reduces chimeric alignments not supported by cDNA sequencing across all cell lines. (i) Chimeric alignments from dRNA-seq of the WTC11 cell line, evaluated for support using additional ONT and Pacific Biosciences (PacBio) sequencing data with different protocols. DeepChopper reduces unsupported chimeric alignments across all methods compared to Dorado with adapter trimming.

To accomplish this, DeepChopper integrates the long-context genomic language model (LCGLM) HyenaDNA [6], optimized for efficient long-range dependency modeling, with a residual block and a multilayer perceptron (MLP) for fine-grained sequence analysis. This hybrid architecture enables both broad context understanding and precise single-nucleotide resolution [7, 8] (Fig. 1a). Additionally, DeepChopper incorporates base quality score information to improve prediction and employs an adaptive tokenization strategy that preserves single-nucleotide resolution while capturing higher-order sequence patterns (See Methods for details). DeepChopper offers several advantages over existing GLMs.

Unlike DNABERT2 [9] and Nucleotide Transformer [10], which are limited to context length and lack single-nucleotide tokenization, DeepChopper processes up to 32 kilobase pairs, a sufficiently long context to capture most mRNAs and can accurately identify non-human reference sequences, such as ONT adapters, with base-pair precision. Moreover, its efficient architecture—using only 7 million parameters—makes DeepChopper computationally practical for large-scale dRNA-seq analysis, in contrast to resource-intensive models like Evo [11], which rely on billions of parameters and may not be feasible in environments with limited computational resources. This combination of long context processing, high-resolution accuracy, and computational efficiency makes DeepChopper particularly well-suited for analyzing massive long-read sequences.

To train DeepChopper for identifying adapter sequences within dRNA-seq long reads, we utilized data from six human cell lines: HEK293T, A549, HCT116, HepG2, K562 and MCF-7 provided by the Singapore Nanopore Expression Project (SG-NEx) [12]. We curated a training set of 540,000 long reads initially deemed free of adapters and inserted putative adapter sequences, derived from the raw dRNA-seq data, into these reads to create instances containing internal and 3′ end adapters (See Methods for details). An independent test set comprising 60,000 long reads was held out for performance evaluation. DeepChopper achieved recall, precision, and F1 scores above 0.99 (Extended Data Fig. 1a), demonstrating its high accuracy in detecting adapter sequences. To further assess DeepChopper’s ability to detect chimera artifacts in real data, we generated an additional dRNA-seq dataset using the prostate cancer VCaP cell line, which was not included in the training data. This independent dataset provides a robust framework for evaluating chimera artifacts in genuine dRNA-seq samples, ensuring that DeepChopper’s performance generalizes beyond the training data.

We conducted dRNA-seq of VCaP cells using ONT‘s SQK-RNA002 chemistry, consistent with that used in the SG-NEx project. Using a MinION sequencer with four R9.4 flow cells, we generated a total of 9,177,639 long reads in FASTQ format, with base-calling performed by ONT‘s Dorado software [13]. DeepChopper was then applied for adapter trimming and correction of chimeric read artifacts. Notably, DeepChopper increased the yield of long-read sequences by 3%, resulting in a final output of 9,357,913 long reads. DeepChopper identified 8,218,172 adapter sequences within 7,990,102 long reads, accounting for 87% of the raw reads. Most of these adapter sequences were approximately 70 bp in length (Fig. 1b), aligning with the expected length of the RMX adapter used in ONT SQK-RNA002 dRNA-seq kit, as specified in the kit’s technical documentation [14].

We then analyzed the positions of DeepChopper identified adapter sequences within each long read. Among these, 212,478 reads contained internal adapters, while 7,777,624 reads had adapters at their 3′ ends (Fig. 1c). This reveals that chimeric read artifacts, evidenced by internal adapters, are present in VCaP dRNA-seq data and may reflect a common problem inherent to dRNA-seq long reads. Further examination of adapter-trimmed reads showed that chimera artifacts could arise from the joining of multiple long reads, with the most prevalent type involving two reads connected by an internal adapter to form a single chimeric read (Fig. 1d).

To validate the chimeric read artifacts identified by DeepChopper in the VCaP dRNA-seq data, we analyzed the chimeric alignments generated by minimap2 [15], as chimeric reads typically produce such alignments. To distinguish genuine chimeric reads from artifacts, we generated nanopore direct cDNA sequencing data for VcaP cells and considered chimeric reads with alignments fully supported by cDNA sequencing as bona fide (See Methods for details). Additionally, we assessed whether the adapter trimming function of ONT‘s Dorado basecaller could mitigate chimeric read artifacts. While Dorado’s adapter trimming had no effect on reducing chimeric alignments, DeepChopper achieved a 91% reduction and increased the proportion of cDNA-supported chimeric alignments from 5% to 47% (Fig. 1e). To further confirm the nature of the identified chimeric read artifacts, we examined the base quality scores of the internal adapter sequences identified by DeepChopper and aligned them with the human reference genome using BLAT [16], a tool known for its high alignment accuracy. Our analysis revealed that the adapter sequences from the chimeric read artifacts exhibited low base quality scores (Fig. 1f) and poor alignment identity with the human reference genome (Fig. 1g), confirming their artifactual nature and non-human origin.

To comprehensively assess DeepChopper’s performance in reducing chimera artifacts in dRNA-seq, we examined cDNA-supported chimeric alignments in the SG-NEx samples. DeepChopper significantly reduced chimeric alignments by 62% to 84%, while preserving cDNA-supported chimeric alignments without noticeable reduction (Fig. 1h), indicating the prevalence of chimeric read artifacts in dRNA-seq.

To further validate DeepChopper’s accuracy, we applied it to process dRNA-seq data from the human WTC11 induced pluripotent stem cell line, generated by the Long-read RNA-Seq Genome Annotation Assessment Project (LRGASP) project [17]. Additionally, the LRGASP project generated cDNA-based long-read sequencing data from this cell line using distinct protocols (PCR-cDNA, CapTrap, R2C2) on both ONT and PacBio platforms, providing a valuable resource for benchmarking DeepChopper’s performance. We observed that DeepChopper consistently eliminated only the chimeric alignments in the dRNA-seq data that were not supported by any cDNA-based sequencing method (Fig. 1i), demonstrating its precise ability to distinguish genuine chimeric reads from artifacts.

Recently, ONT released a new SQK-RNA004 chemistry for dRNA-seq, but it remains unclear whether chimera artifacts persist with this update. To investigate, we generated dRNA-seq data using the latest SQK-RNA004 kit for the VCaP cell line and performed zero-shot predictions of chimera artifacts using DeepChopper, despite the adapter sequence pattern of RNA004 not being included in its training. DeepChopper reduced chimeric alignments by 21% compared to Dorado’s base-called and adapter-trimmed long reads, increasing the proportion of cDNA-supported chimeric alignments from about 25% to 30% (Extended Data Fig. 1b). Furthermore, the internal adapter sequences identified by DeepChopper within these RNA004 chimeric read artifacts exhibited low base quality scores (with a mean of 7.8) and low BLAT identity (mean of 0.38) (Extended Data Fig. 1c). These findings strongly suggest that chimera artifacts continue to be an issue with the latest RNA004 dRNA-seq kit.

To investigate potential factors contributing to the formation of chimera artifacts, we examined the expression levels and sizes of the genes involved in these artifacts, we found that genes affected by chimera artifacts exhibited significantly higher expression levels compared to all expressed genes (p-value *<* 2.2 *×* 10^−16^) (Extended Data Fig. 2a), while following a similar size distribution pattern (Extended Data Fig. 2b). We then analyzed the genomic patterns of improper connections generated by chimera artifacts across chromosomes in VCaP dRNA-seq data. These artifacts showed varying frequencies across chromosomes, with the mitochondrial chromosome (Chr M) exhibiting an especially high rate of connections per base pair, indicating it may be a hotspot for chimera artifact formation (Fig. 2a). A similar pattern of improper inter-chromosomal connections was also observed in VCaP RNA004 dRNA-seq data (Extended Data Fig. 2c), suggesting that chimera artifacts may be an inherent issue of dRNA-seq technology, rather than being fully resolved by improvements in sequencing chemistry.

**Fig. 2.**
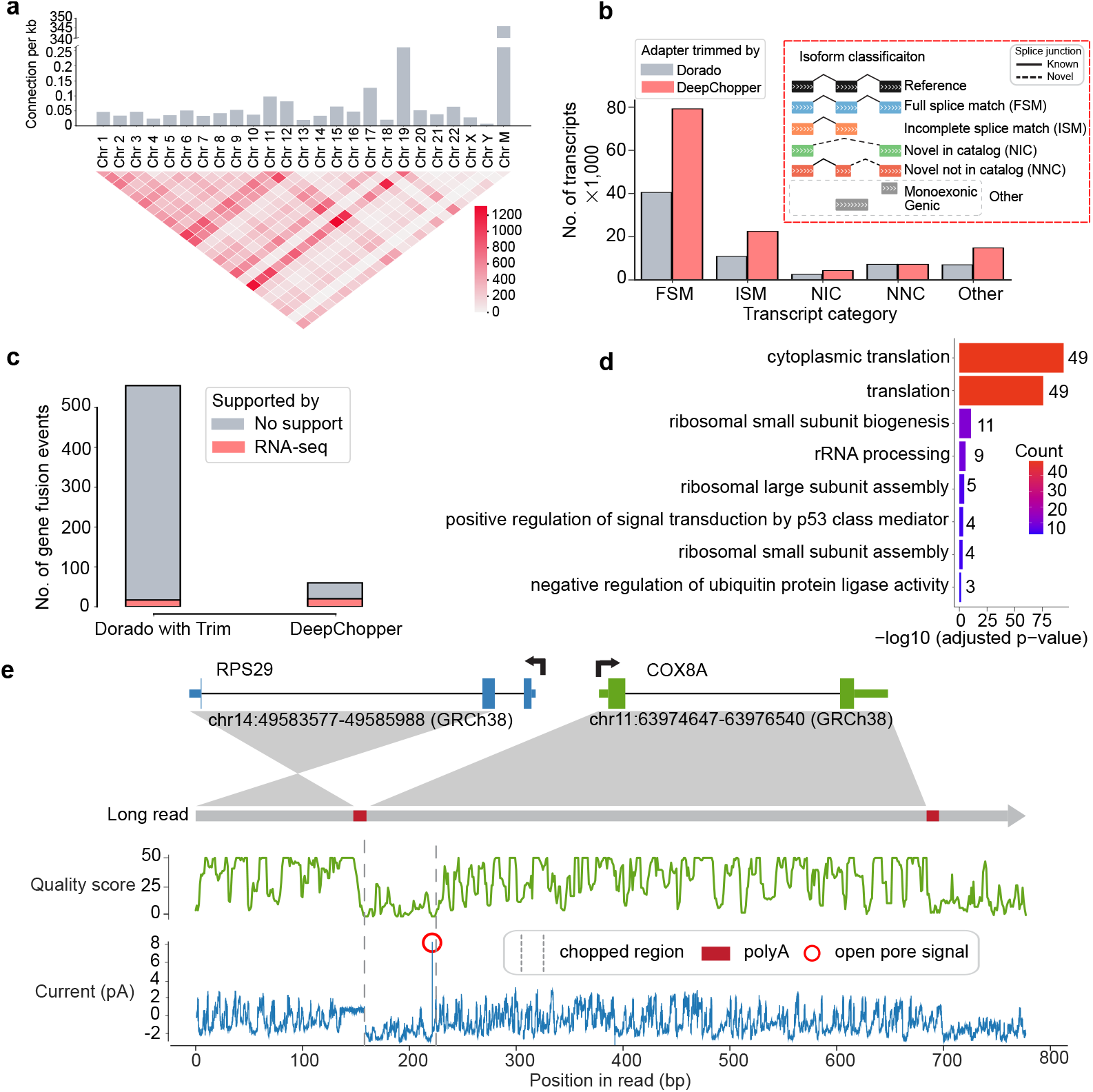
Characterization of dRNA-seq chimera artifacts and their impact on downstream analysis in VCaP cells. (a) The upper bar plot shows the number of chimeric connections per kilobase across chromosomes, highlighting higher chimeric activity in Chr 19 and Chr M. The lower heatmap visualizes interchromosomal connections, with intensity indicating the count of connections between different chromosomes. (b) The bar plot shows the number of transcripts (in thousands) across different isoform classification categories. DeepChopper-processed reads result in a higher number of transcripts compared to Dorado-trimmed reads. The inset details the isoform classification scheme. (c) Detected gene fusions from Dorado adapter-trimmed reads and DeepChopper-processed reads. Gene fusions identified from short-read RNA-seq were used to validate fusion events detected from dRNA-seq. (d) Gene Ontology (GO) enrichment analysis of chimera artifact-affected genes, with color indicating gene count per term. (e) Analysis of a chimeric read artifact detected as an RPS29-COX8A fusion. The schematic shows the fusion between RPS29 (Chr 14) and COX8A (Chr 11). The green plot indicates quality scores along the read, and the blue plot shows raw signal intensity (in pA). The chopped region identified by DeepChopper corresponds to a low-quality segment with low current intensity, polyA, and short open pore signals, suggesting the presence of an ONT adapter.

To evaluate the influence of chimera artifacts on downstream analyses, we investigated the impact of DeepChopper-mediated correction on transcript annotation. IsoQuant [18] was employed to detect and annotate transcripts from VCaP dRNA-seq data, focusing on chimeric read artifacts identified by DeepChopper. By comparing transcript identification with and without DeepChopper correction, we observed an approximately 2-fold increase in the number of identified transcripts (Fig. 2b). Similar results were also noted for VCaP RNA004 dRNA-seq data (Extended Data Fig. 2d). The most prominent increase occurred in the full-length transcripts (Full splice match (FSM) category), with additional improvements observed across alternatively spliced transcripts (Incomplete splice match (ISM), Novel in catalog (NIC), and Novel not in catalog (NNC) categories). These findings underscore the effectiveness of DeepChopper in mitigating the detrimental effects of chimera artifacts on transcript annotation.

Next, we assessed whether DeepChopper could improve gene fusion detection by correcting chimera artifacts. Gene fusion calls identified by FusionSeeker [19] from DeepChopper-processed dRNA-seq reads exhibited an 89% reduction compared to those detected from Dorado’s adapter-trimmed reads. To further evaluate this reduction, we applied Arriba [20] to detect gene fusions using short-read RNA-seq of VCaP cells and found that none of the gene fusion calls reduced by DeepChopper were supported by those detected in RNA-seq data (Fig. 2c). This demonstrates that DeepChopper effectively reduces false positive gene fusion calls caused by chimera artifacts in dRNA-seq, an essential improvement for accurately identifying gene fusion drivers in cancer transcriptomes.

Upon closer examination of the chimera artifact-derived gene fusion calls, we found a notable enrichment of ribosomal protein genes (Extended Data Fig. 3a). GO enrichment analysis in VCaP (Fig. 2d) and SG-NEx cell lines (Extended Data Fig. 3b) confirmed that housekeeping ribosomal protein genes are frequently involved in these artifacts. Additionally, ribosomal protein genes were also enriched in the chimera artifacts associated with RNA004 chemistry (Extended Data Fig. 3b). A manual inspection of a chimeric read identified as an RPS29-COX8A fusion, involving the 40S ribosomal protein S29 in VCaP dRNA-seq data, revealed that the DeepChopper-processed region aligned with raw current signals showing low intensity, indicative of ONT adapters (Fig. 2e). The presence of polyA and open pore signals at the boundary of this region further suggested that this chimeric read resulted from improperly joined mRNA transcripts, rather than representing a genuine gene fusion event.

In conclusion, DeepChopper demonstrates a significant potential for leveraging LLM to analyze long-read sequences, particularly by addressing the previously overlooked issue of chimera artifacts in nanopore dRNA-seq. DeepChopper significantly enhances the quality of the dRNA-seq data by identifying and removing ONT adapters directly from the base-called long-read sequences, thereby boosting the precision of subsequent transcript and gene fusion analyses. This advancement not only optimizes current nanopore dRNA-seq analysis but also highlights the broader applicability of language models to address the complex biological challenges posed by emerging long-read technologies.

## Methods

### Cell culture

VCaP cell line was obtained from the American Type Culture Collection (ATCC) and cultured under sterile conditions to maintain optimal growth and viability. The cells were grown in Dulbecco’s Modified Eagle Medium (DMEM, high glucose; Gibco, Cat# 11-965-092) supplemented with 10% fetal bovine serum (FBS Opti-Gold, Performance Enhanced, US Origin; Gendepot, Cat# F0900-050) to provide essential growth factors. In addition, the culture medium was enriched with 5 mL of 100 mM Sodium Pyruvate (Gendepot, Cat# CA017-010) to support cellular metabolism and 5 mL of Antibiotics-Antimycotics (100×) (Gendepot, Cat# CA002-010) to prevent microbial contamination. Cells were cultured in 100 mm cell culture treated dishes (Thermo Fisher Scientific, Cat# 12-556-002) and incubated at 37 °C in a humidified atmosphere containing 5% CO2, with media changes performed every 72 hours to ensure nutrient availability and waste removal. Cell confluency was regularly monitored and subculturing was performed before reaching 80% confluency to maintain healthy growth conditions and prevent over-confluence stress.

### RNA extraction and quantification

Total RNA was extracted using the RNeasy Mini Kit (Qiagen, Cat# 74104) according to the protocol of the manufacturer. The quality and concentration of RNA were assessed using an Agilent 2100 Bioanalyzer. Poly(A)+ RNA was then enriched from total RNA using the Dynabeads^tm^ mRNA Purification Kit (Invitrogen, Cat# 65001), which utilizes oligo (dT) beads for selective mRNA binding. The mRNA was quantified using a Qubit 4 fluorometer and a Qubit RNA HS Assay Kit (Thermo Fisher Scientific, Cat# Q32852). The mRNA preparations were either immediately used to prepare a sequencing library or frozen and stored at −80 °C until further use.

### Nanopore sequencing

We performed nanopore dRNA-seq sequencing of the enriched mRNA using two different sets: the RNA002 kits with R9.4.1 flow cells and the RNA004 kits with R10.4.1 flow cells. The decision to incorporate the RNA004 kit, a newly released option, was driven by our intention to test its capabilities in conjunction with our DeepChopper tool to optimize data quality and sequencing efficiency. For the RNA002 library, 1 μg of poly(A)+ RNA was used as input for library preparation using the Direct RNA Sequencing Kit (SQK-RNA002, ONT) following the manufacturer’s instructions. Nanopore dRNA-seq employs a reverse transcriptase adapter (RTA) that typically binds to the poly(A) tails of messenger RNA (mRNA); subsequently, a sequencing adapter is ligated to the RTA, which guides the mRNA through the nanopore for sequencing. The prepared library was loaded onto four MinION R9.4 flow cells (FLO-MIN106) and sequenced for 48 hours using the Oxford Nanopore MinION device. For the RNA004 library, 300 ng of poly(A)+ RNA was used as input for library preparation using the Direct RNA Sequencing Kit (SQK-RNA004, ONT) according to the protocol of the manufacturer. The library was then loaded onto a PromethION RNA Flow Cell (FLO-PRO004RA) and sequenced on the Oxford Nanopore PromethION device for 72 hours.

For Direct cDNA sequencing, we utilized the Direct cDNA Sequencing Kit (SQK-DCS109, ONT) following the manufacturer’s protocol. Briefly, 5 μg of total RNA was used as input for first-strand cDNA synthesis using Maxima H Minus Reverse Transcriptase (Thermo Fisher Scientific) with the SSP and VN primers provided in the kit. To eliminate potential RNA contamination, we treated the sample with RNase Cocktail Enzyme Mix (Thermo Fisher Scientific). Second-strand cDNA synthesis was carried out using LongAmp Taq Master Mix (New England Biolabs). The resulting double-stranded cDNA underwent end-repair and dA-tailing using the NEBNext Ultra End Repair/dA-Tailing Module (New England Biolabs). Subsequently, sequencing adapters were ligated to the prepared cDNA using Blunt/TA Ligase Master Mix (New England Biolabs). Between each enzymatic step, the cDNA and libraries were purified using AMPure XP beads (Agencourt, Beckman Coulter). We quantified the libraries using a Qubit Fluorometer 3.0 (Life Technologies) to ensure adequate concentration and quality. The final library was loaded onto a MinION R9.4 flow cell and sequenced on the Oxford Nanopore MinION device for 72 hours.

### Training data preparation

We acquired ONT dRNA-seq FAST5 data from the SG-NEx project, which includes six human cell lines: HEK293T, A549, K562, HepG2, MCF7, and HCT116 [12]. The FAST5 files were converted to POD5 format using the POD5 conversion tool (https://pod5-file-format.readthedocs.io). Subsequently, FASTQ files were generated using Dorado (v0.5.2) [13] with adapter trimming disabled (`--no-trim`) and the “rna002 70bps hac@v3” model. The reads were then aligned to the human reference genome (GRCh38) using minimap2 (v2.24) [15] with ONT direct RNA-specific parameters (-ax splice -uf -k14) for optimized alignment. The resulting SAM files were then converted to BAM format, indexed, and sorted using SAMtools (v1.19.2) [21]. For adapter sequence extraction, we selected primary alignments without supplementary alignments and identified 3′ end soft-clipped regions as adapter sequences, while aligned regions were considered non-adapter sequences. To create artificial chimeric reads, we randomly combined two non-adapter sequences with one adapter sequence to create FASTQ records. The dataset consists of positive examples containing adapter sequences (with a 1:1 ratio of 3′ end and internal adapters) and negative examples without any adapter sequences, in a 9:1 ratio. In total, 600,000 data points were generated and divided into training (480,000), validation (60,000), and test sets (60,000) in an 8:1:1 ratio using stratified random sampling.

### Language model architecture

DeepChopper identifies adapter sequences by framing the problem as a token classification task. The model tokenizes biological sequences at the single-nucleotide level, treating each nucleotide (*A, C, G, T*, and *N*) as a fundamental unit. This approach enables fine-grained sequence analysis crucial for distinguishing artificial adapter sequences from native biological sequences.

For each nucleotide token, DeepChopper predicts whether it belongs to an adapter sequence. The model’s architecture is optimized for high-precision identification of adapter sequences within biological data. At its core, DeepChopper utilizes the Hye-naDNA [6], a specialized genomics architecture that serves as the primary feature extractor. This framework efficiently processes nucleotide sequences to produce rich feature representations with a dimensionality of 256.

These extracted features then pass through a quality block that integrates base quality scores. The quality block consists of dense layers with residual connections, incorporating sequence quality information to enhance the model’s interpretation of the input data.

The enriched features are ultimately fed into a classification head that computes the probability of each token being part of an adapter sequence. Through binary classification at every nucleotide position, the model labels each token as either adapter or non-adapter sequence, providing a comprehensive analysis of the entire input sequence.

### Model training

DeepChopper processes sequences up to 32,770 nucleotides in length, excluding any longer sequences from analysis. To ensure efficient batch processing, shorter sequences were padded to this maximum length. The model was trained using a supervised learning approach, utilizing sequences labeled with adapter annotations. Training was performed in a High Performance Computing (HPC) cluster using two A100 Graphics Processing Units (GPUs). The batch size was set to 64, and validation was performed every 20,000 steps. The model with the highest validation F1 score for the base prediction task was selected for subsequent analyses. Training was carried out over 60 epochs, with early stopping applied based on validation performance to mitigate overfitting risks.

The Adam optimizer was used for parameter optimization, with settings of *β*_1_ = 0.9 and *β*_2_ = 0.999 [22]. A learning rate scheduler was used to reduce the learning rate when validation loss ceased improving, starting with an initial learning rate of 2 × 10^−5^. The cross-entropy loss function was used to update the model parameters, defined as follows:

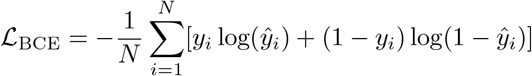

where ℒ_*BCE*_ is the binary cross-entropy loss, *N* is the total number of tokens in the input sequence, *y*_*i*_ is the ground true label for the *i*-th token, and *ŷ*_*i*_ is the predicted probability for the *i*-th token.

The average cross-entropy loss across the mini-batch is computed as:

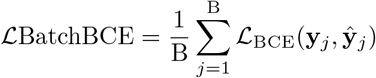

where ℒ_*BatchBCE*_ is the average binary cross-entropy loss for the mini-batch, B is the batch size (number of sequences in the mini-batch), and **y**_*j*_ and **ŷ**_*j*_ are the true labels and predicted probabilities for the *j*-th sequence in the batch.

The model evaluation metrics included accuracy, precision, recall and the F1 score, calculated using the following equations:

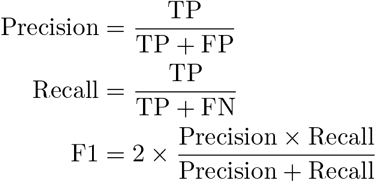

The final selection of the model was based on the optimal performance in the validation set. The model is implemented by PyTorch (v2.5.0) [23]. To identify the best hyperparameter configuration, the Hydra (v1.3.2) [24] framework was used.

### Sliding window approach for prediction refinement

To improve the accuracy and smoothness of the predictions, a sliding window approach was implemented. This method extends the predicted regions and minimizes noisy predictions, aligning with the typical length of adapter sequences observed in the ONT dRNA-seq data. The sliding window strategy effectively maintains continuity in regions predicted to contain adapter sequences, preventing fragmented and unrealistic results.

A sliding window size of 21 nucleotides was selected based on empirical optimization, balancing the need for smoothing while retaining sensitivity to shorter adapter sequences. Within each window, a voting mechanism was applied to refine the predictions. The final classification *y*_*i*_ for each nucleotide *x*_*i*_ was determined by the majority vote of all predictions within the window, as defined by the following equation:

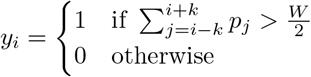

where *y*_*i*_ is the final prediction for the *i*-th nucleotide, *W* is the sliding window size, *k* is half the window size 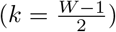, and *p*_*j*_ represents the initial predicted label for the *j*-th nucleotide within the window, where a value of 1 indicates that the nucleotide is part of an adapter sequence, and a value of 0 indicates that it is part of a non-adapter sequence.

The combination of the sliding window technique and voting-based refinement significantly improved prediction accuracy by smoothing the outputs and reducing false positives, resulting in more reliable data essential for downstream analysis.

### Post-processing and filtering

After refining the adapter predictions, four filtering steps were applied to enhance the quality of the final results:

1. A predicted adapter sequence must be at least 13 nucleotides long. Sequences shorter than this length threshold are not considered valid adapters.
2. If a read contains more than four adapter sequences, the entire read sequence is retained without any adapter removal.
3. For reads containing four or fewer adapter sequences, the identified adapters are removed and the read is divided into smaller segments.
4. Any segments resulting from this process that are less than 20 nucleotides long are discarded.

Each remaining segment and its corresponding base quality scores are stored as a single read record in the final FASTQ file. This filtering process separates chimeric read artifacts containing internal adapters into multiple segments, while retaining reads with 3′ end adapters as single shortened segments.

### BLAT identity calculation

The accuracy of DeepChopper in detecting adapter sequences was evaluated by aligning the identified sequences to the human reference genome using BLAT [16]. A BLAT identity score was defined as the ratio of matched bases to the total sequence length:

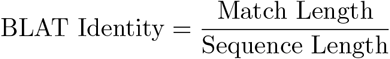

In this context, match length refers to the number of bases in the query sequence that align with the reference genome, while sequence length denotes the total length of the query sequence. This score provides a quantitative measure of how closely each identified sequence aligns with the reference genome, serving as an indicator of detection accuracy. The alignments were performed using the PxBLAT (v1.2.1) [25]

### Cross-platform validation

Cross-platform validation of dRNA-seq chimera artifacts identified by DeepChopper was conducted leveraging ONT direct cDNA sequencing and additional cDNA-based sequencing platforms. Direct cDNA sequencing validation was performed using six cancer cell lines, including the VCaP dataset generated in this study and five published datasets (A549, K562, HepG2, MCF7, and HCT116) obtained from the SG-NEx project [12]. The direct cDNA data in FAST5 format were converted to POD5 format using the POD5 conversion tool (https://pod5-file-format.readthedocs.io). Subsequently, FASTQ files were generated using Dorado (v0.5.2) [13] with adapter trimming enabled (--trim adapters) and the “dna r9.4.1 e8 hac@v3.3” model. The reads were then processed using Pychopper (https://github.com/epi2me-labs/pychopper, v2.7.9) and Cutadapt (v4.2) [26] according to a published protocol [27]. The oriented reads were aligned to the human reference genome (GRCh38) using minimap2 (v2.24) [15] with optimized parameters (-ax splice -uf -k14) for spliced alignment. The resulting SAM files were then converted to BAM format, indexed, and sorted using SAMtools (v1.19.2) [21].

Additional cDNA-based long-read sequencing data from the WTC11 cell line were used for further validation, incorporating five distinct platforms: ONT PCR-cDNA, ONT CapTrap, ONT R2C2, PacBio cDNA, and PacBio CapTrap. The raw FASTQ files (and FASTA files for ONT R2C2) from these datasets were provided by the LRGASP project [17]. For the PCR-cDNA data, the reads were processed using Pychopper (https://github.com/epi2me-labs/pychopper, v2.7.9) and Cutadapt (v4.2) [26], following the protocol described in reference [27]. ONT reads were then aligned to the human reference genome (GRCh38) using minimap2 (v2.24) [15] with the parameters (-ax splice -uf -k14), while PacBio reads were aligned using the parameters (-ax splice:hq -uf). The ONT dRNA-seq data from A549, K562, HepG2, MCF7, HCT116, VCaP, and WTC11 cell lines were processed as previously described.

To validate the chimeric alignments derived from dRNA-seq, comparisons were made with chimeric alignments identified from cDNA-based data across the specified platforms. Chimeric alignments, defined by a primary alignment and one or more supplementary alignments, each containing the SA tag in the BAM file, were converted into lists of genomic intervals based on their corresponding alignments. The genomic interval lists were then compared between platforms, and overlapping intervals were considered concordant if the distance difference between them was less than 1000 bp. Supporting rates were calculated as the proportion of dRNA-seq chimeric alignments corroborated by cDNA-based platforms, thereby providing cross-validation of chimera artifacts identified by DeepChopper.

### analysis and transcript classification

Gene expression levels from dRNA-seq were quantified using IsoQuant (v3.1.2) [18], with the parameters (--data type nanopore --stranded forward --model construction strategy default ont --sqanti output). The “--sqanti output” option enables IsoQuant to generate files containing transcript classification information, analogous to the output provided by SQANTI [28].

### Gene fusion identification and visualization

For ONT dRNA-seq data, gene fusions were identified using FusionSeeker (v1.0.1) [19] with default settings. For short-read RNA-seq data, FASTQ files for the VCaP cell line were obtained from the Cancer Cell Line Encyclopedia (CCLE) project [29] under SRA accession SRX5417211. Raw reads were mapped to the hg38 reference genome using STAR (v2.7.11) [30], and gene fusion events were detected with Arriba (v2.4.0) [20]. The gene structure of the RPS29-COX8A fusion was visualized using GSDS (v2.0) [31]. Base quality scores were generated with a custom Python script, and ion current signals were visualized using Squigualiser (v0.6.3) [32]. The circos plot for gene fusion events was visualized using chimeraviz (v1.30.0) [33].

### GO enrichment analysis

GO enrichment analysis of biological processes for genes involved in chimera artifacts identified in dRNA-seq data was performed using the Database for Annotation, Visualization, and Integrated Discovery (DAVID) webserver [34].

### Computing resource

All computations were performed on a HPC server equipped with a 64-core Intel(R) Xeon(R) Gold 6338 CPU and 256 GB of RAM. The server was also configured with two NVIDIA A100 GPUs, each with 80 GB of memory, enabling efficient processing of both CPU-intensive tasks and GPU-accelerated deep learning workloads.

## Data Availability

Raw and processed data generated in this study, including dRNA-seq using the SQK-RNA002 and SQK-RNA004 kits, as well as direct cDNA sequencing of VCaP cells, have been deposited in the Gene Expression Omnibus (GEO) under the accession number GSE277934.

## Code Availability

DeepChopper, implemented in Rust and Python, is open source and available on GitHub (https://github.com/ylab-hi/DeepChopper) under the Apache License, Version 2.0. The package can be installed via PyPI (https://pypi.org/project/deepchopper/) using pip, with wheel distributions provided for Windows, Linux, and macOS to ensure easy cross-platform installation. An interactive demo is available on Hugging Face (https://huggingface.co/spaces/yangliz5/deepchopper), allowing users to test DeepChopper’s functionality without local installation. For large-scale analyses, we recommend using DeepChopper on systems with GPU acceleration. Detailed system requirements and optimization guidelines are available in the repository’s documentation.

## Acknowledgements

This project was supported in part by NIH grants R35GM142441 and R01CA259388 awarded to RY, and NIH grants R01CA256741, R01CA278832, and R01CA285684 awarded to QC.

## Author Contributions

YL, TYW and RY designed the study with QC. YL and TYW performed the analysis. QG, YR and XL performed the experiments. YL designed and implemented the model and computational tool. YL, TYW, QG and RY wrote the manuscript. RY supervised this work.

## Acronyms

This document is incomplete. The external file associated with the glossary ‘acronym’ (which should be called `output.acr`) hasn’t been created.

Check the contents of the file `output.acn`. If it’s empty, that means you haven’t indexed any of your entries in this glossary (using commands like \gls or \glsadd) so this list can’t be generated. If the file isn’t empty, the document build process hasn’t been completed.

Try one of the following:

- Add `automake` to your package option list when you load `glossaries-extra.sty`.For example: `\usepackage[automake]{glossaries-extra`}
- Run the external (Lua) application: `makeglossaries-lite.lua “output”`
- Run the external (Perl) application: `makeglossaries “output”`

Then rerun LATEX on this document.

This message will be removed once the problem has been fixed.

## Extended data

**Extended Data Fig. 1.**
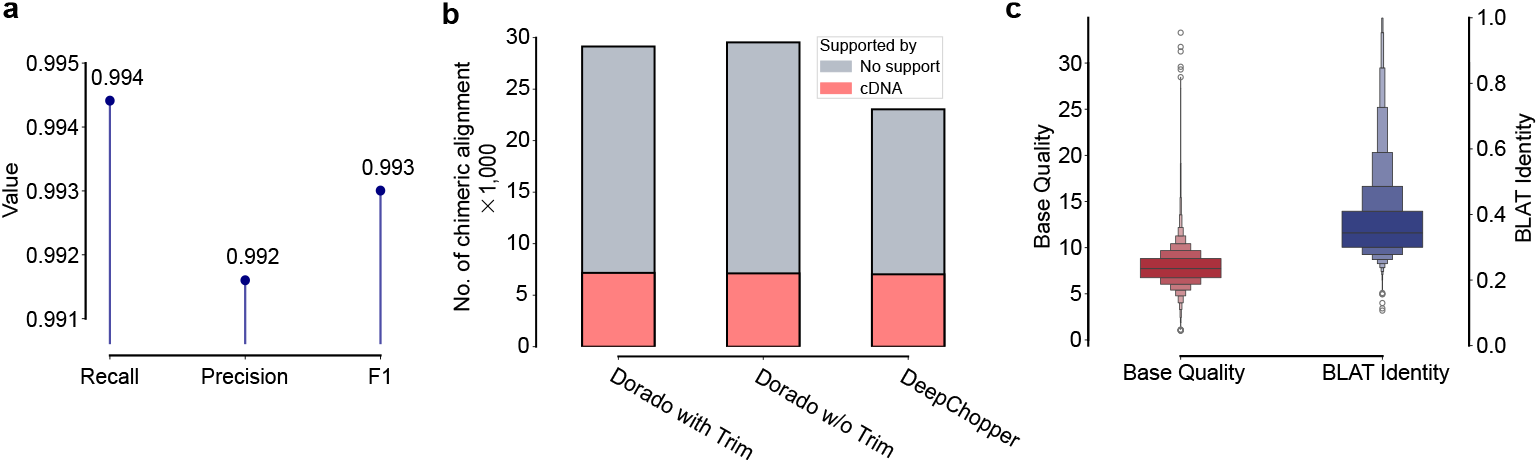
Benchmarking of DeepChopper predictions on chimeric read artifacts using synthetic data and dRNA-seq data generated with the SQK-RNA004 kit from the VCaP cell line. (a) Performance evaluation in a held-out test dataset showing Recall, Precision, and F1 values for DeepChopper. (b) Number of chimeric alignments (in thousands) identified in VCaP RNA004 dRNA-seq reads processed by Dorado with and without adapter trimming, and by DeepChopper. The bar colors indicate chimeric reads supported by cDNA sequencing (red) and those lacking support (grey). (c) Base quality scores (left) and BLAT alignment identity scores (right) for internal adapter sequences identified by DeepChopper in RNA004 dRNA-seq reads.

**Extended Data Fig. 2.**
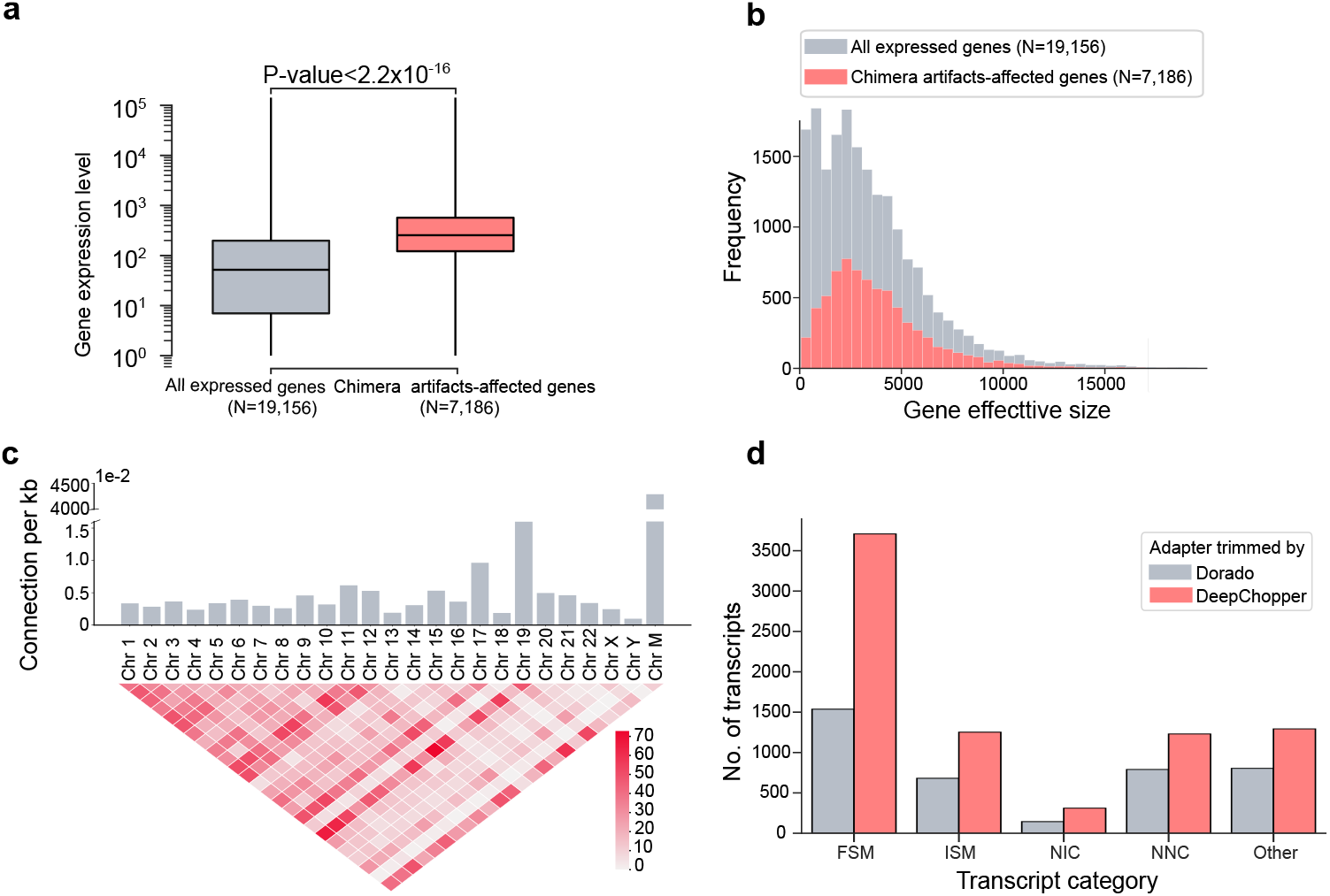
Analysis of dRNA-seq chimera artifacts and their genomic and transcriptomic characteristics in VCaP cells. (a) Box plot comparing gene expression levels between all expressed genes (N=19,156) and genes affected by chimera artifacts (N=7,186) in the VCaP dRNA-seq dataset. Chimera artifacts-affected genes exhibit significantly higher expression levels (p-value *<* 2.2 × 10^−16^). (b) Distribution of gene effective sizes for all expressed genes and genes affected by chimera artifacts, indicating that the size distributions of genes impacted by chimera artifacts are comparable to those of all expressed genes. (c) Chromosomal distribution and interchromosomal connections from chimeric read artifacts arising from VCaP RNA004 dRNA-seq. The top bar plot shows the number of connections per kilobase for each chromosome, with higher bars indicating more frequent connections. The bottom heatmap visualizes the number of chimeric connections between chromosome pairs, with color intensity representing the connection frequency. (d) Number of detected transcripts across different isoform categories (FSM, ISM, NIC, NNC, and Other) from DeepChopper-identified chimeric read artifacts in VCaP RNA004 dRNA-seq data. DeepChoppercorrected reads resulted in a greater number of transcripts compared to adapter-trimmed reads by Dorado across all categories.

**Extended Data Fig. 3.**
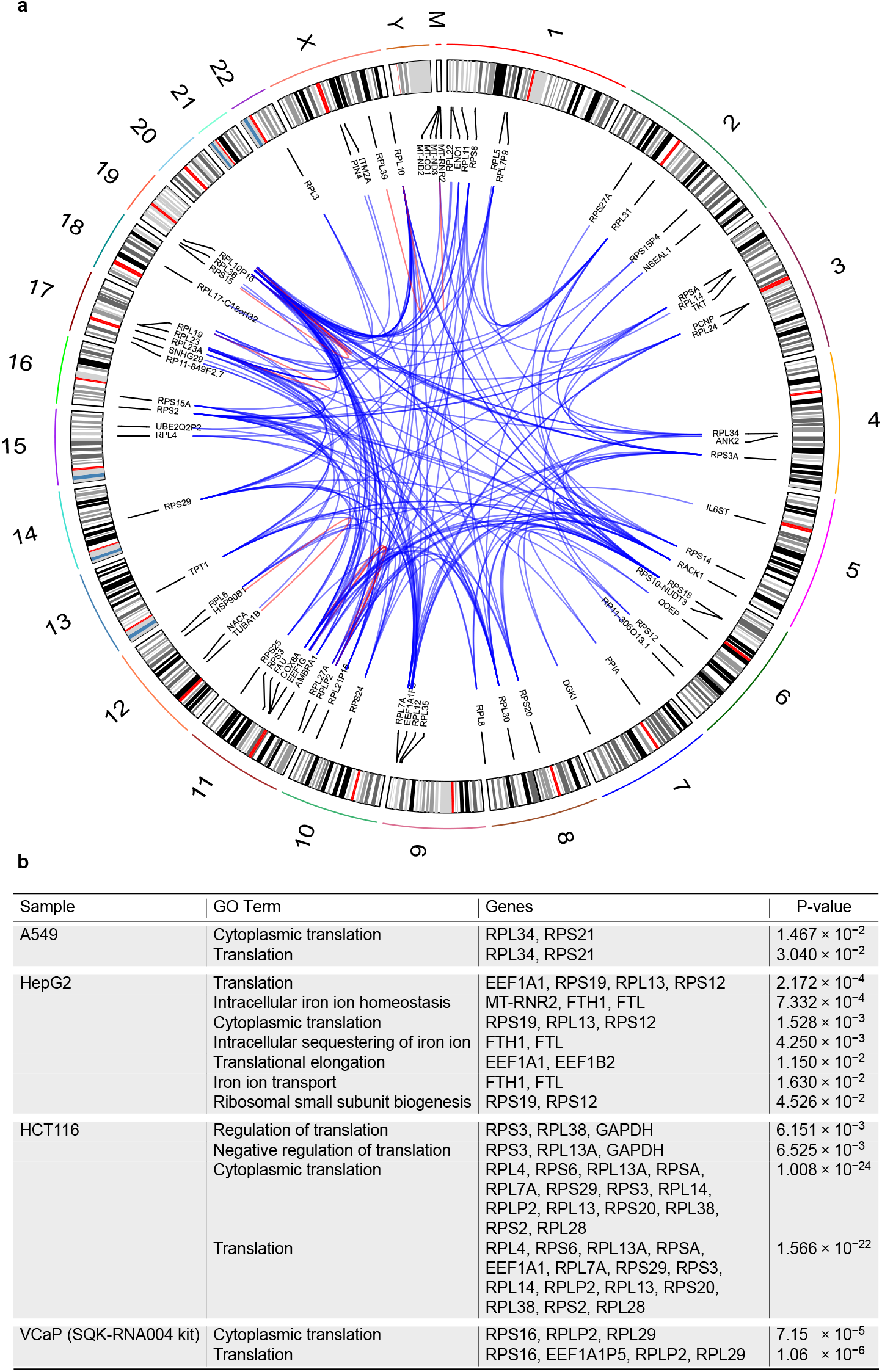
Analysis of gene fusions derived from chimeric read artifacts in dRNA-seq. (a) Circos plot depicting chromosomal connections of gene fusions resulting from chimeric read artifacts in VCaP cells. Blue lines represent inter-chromosomal fusion events, while red lines indicate intra-chromosomal fusions. The outer track displays chromosomal ideograms labeled with respective chromosome numbers. (b) GO enrichment analysis of fusion genes derived from chimeric read artifacts identified by DeepChopper in dRNA-seq data from A549, HepG2, and HCT116 cell lines, and VCaP RNA004 dRNA-seq data. The table lists enriched GO terms of biological processes, associated genes, and the statistical significance (p-values) for each enrichment.

